# The role of conformational entropy in the determination of structural-kinetic relationships for helix-coil transitions

**DOI:** 10.1101/237875

**Authors:** Joseph F. Rudzinski, Tristan Bereau

## Abstract

Coarse-grained molecular simulation models can provide significant insight into the complex behavior of protein systems, but suffer from an inherently distorted description of dynamical properties. We recently demonstrated that, for a heptapeptide of alanine residues, the structural and kinetic properties of a simulation model are linked in a rather simple way, given a certain level of physics present in the model. In this work, we extend these findings to a longer peptide, for which the representation of configuration space in terms of a full enumeration of sequences of helical/coil states along the peptide backbone is impractical. We verify the structural-kinetic relationships by scanning the parameter space of a simple native-biased model and then employ a distinct transferable model to validate and generalize the conclusions. Our results further demonstrate the validity of the previous findings, while clarifying the role of conformational entropy in the determination of the structural-kinetic relationships. More specifically, while the global, long timescale kinetic properties of a particular class of models with varying energetic parameters but approximately fixed conformational entropy are determined by the overarching structural features of the ensemble, a shift in these kinetic observables occurs for models with a distinct representation of steric interactions. At the same time, the relationship between structure and more local, faster kinetic properties is not affected by varying the conformational entropy of the model.

## I. INTRODUCTION

Coarse-grained (CG) molecular simulation models have played a key role in laying the foundation for modern theories of protein folding, and continue to provide significant insight into the complex dynamical processes sampled by biological macromolecules. ^1, 2^ These models are useful for providing microscopic interpretations that complement experimental findings, especially for systems and processes that are computationally out of reach for atomically-detailed models. For example, single-molecule experiments probe a large range of complex kinetic processes sampled by biomolecular systems, but require an underlying molecular model for accurate structural interpretations.^3^ The computational effort required to investigate a system not only depends on the sheer number of particles but also on the range of relevant timescales, thermodynamic and chemical conditions, as well as system variations (i.e., mutations). Therefore, it is clear that CG models will continue to be essential for providing consistent and exhaustive interpretations for experimental observations.

Despite significant advances in the development of chemically-specific CG models for proteins,^4–7^ a fundamental challenge severely limits the predictive capabilities of CG models—the interpretation of CG dynamical properties. The process of removing degrees of freedom from a system typically results in decreased molecular friction and softer interaction potentials. This effect is a double-edged sword: effectively speeding up the sampling of configuration space while obscuring the connection to the true dynamics of the system. These “lost” dynamics not only prevent quantitative prediction of kinetic properties, but may also lead to qualitatively incorrect interpretations generated from CG simulations.^8, 9^ Unlike many polymer systems, where a homogeneous dynamical rescaling factor is capable of recovering the correct dynamics of the underlying system,^10, 11^ the rescaling associated with the complex hierarchy of dynamical processes generated by biological molecules is likely a complex function of the system’s configuration. Consequently, the application of bottom-up approaches that aim to re-insert the appropriate friction via a generalized Langevin equation remains conceptually and computationally challenging.12

We recently demonstrated that, if a CG model incorporates certain essential physics, simple relationships between structural and kinetic properties may emerge.^13^ These structural-kinetic relationships represent powerful tools that can be employed to ensure a CG protein model generates consistent kinetic information, in terms of both relative timescales of dynamical processes and the dynamical pathways sampled during a particular process. To identify these relationships we considered a model system for helix formation—a heptapeptide of alanine residues—and employed a flavored-Gō model,^14–16^ whose parameters are easily tuned to generate particular structural features. To ensure accurate modeling of the structural ensemble, i.e., to avoid sterically forbidden conformations, we incorporated a detailed representation of steric interactions into the model. We then performed a systematic search through parameter space, afforded by the simplicity of the model, and analyzed correlations between the emergent structural and kinetic properties. To validate the generality of our conclusions, we also considered a transferable model with more complex interaction potentials but a slightly simpler representation of steric interactions.

In this study, we investigate the robustness of our previous conclusions by considering a longer peptide with experimental reference data—the capped, helix forming peptide AC-(AAQAA)_3_ - NH_2_. We follow the strategy of our previous study, while expanding upon the models employed, which clarifies the impact of model representation or, more precisely, conformational entropy on the resulting structural-kinetic relationships. More specifically, by varying the energetic parameters of a given model type, we keep the conformational entropy approximately fixed and demonstrate that the global, long timescale kinetic properties (i.e., ratio of folding to unfolding timescales) are determined precisely by the average helical content of the ensemble. Comparison between two distinct model types demonstrates a shift in the timescale ratios due to the change in model representation. Furthermore, by adjusting the steric interactions of one model type, we provide clear evidence that the conformational entropy is the dominant contributor to this shift. In contrast, we find that more local, faster kinetic processes are consistently determined by structural features of the ensemble, regardless of the precise model representation.

## II. COMPUTATIONAL METHODS

### A. Coarse-grained (CG) models

#### 1. Hybrid Gō (Hy-Gō)

To investigate the relationship between structural and kinetic properties generated from CG simulation models, we employ a Gō-type model,^17^ which defines attractive interactions based on the location of atoms in the native structure. More specifically, we use a flavored-Gō model with three simple, Gō-type parameters:^14–17^ *(i)* a native contact (nc) attraction, *U*_nc_, employed between pairs of *C_α_* atoms which lie within a certain distance in the native structure, i.e., the *α*-helix, of the peptide, *(ii)* a desolvation barrier (db) interaction, *U*_db_, also employed between native contacts, and *(iii)* a hydrophobic (hp) attraction, *U*_hp_, employed between all pairs of C_*β*_ atoms of the amino acid side chains. The same functional forms are employed as in many previous studies,^16^ with a tunable prefactor for each interaction as described below. The form of the interactions are illustrated in the top two panels of Figure 1.

**FIG. 1.**
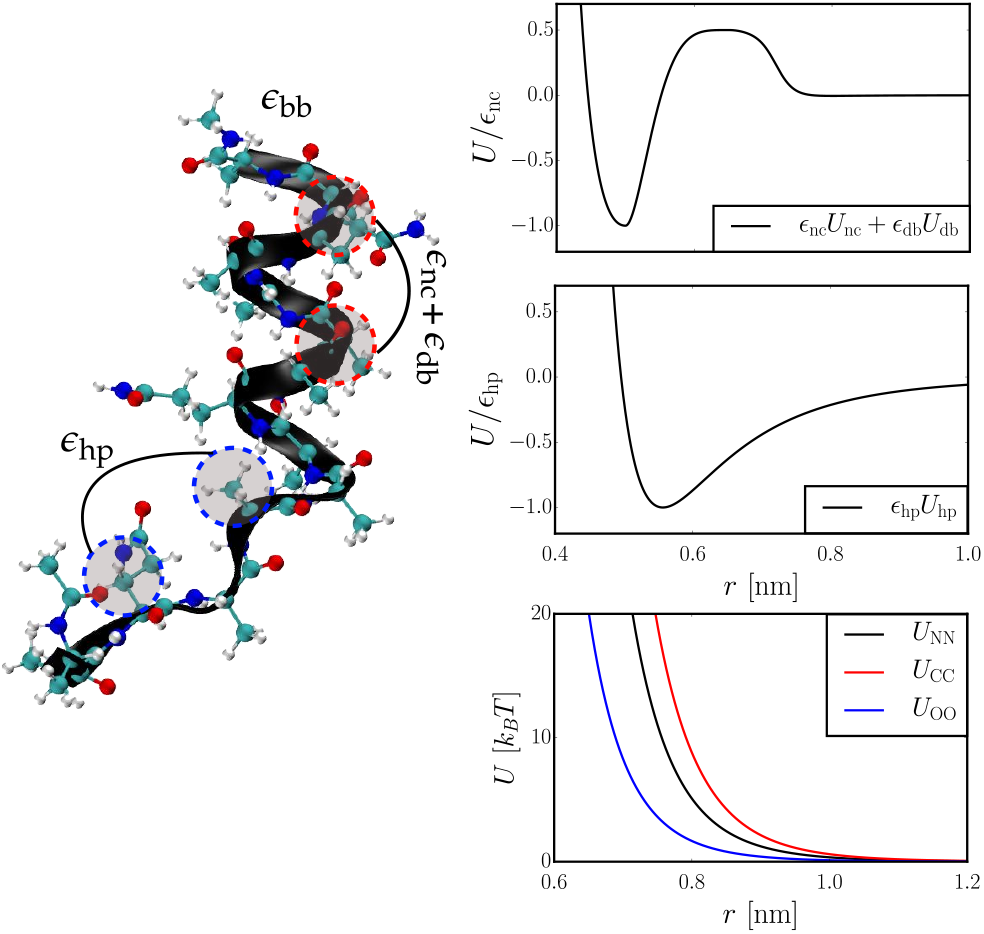
A visualization of the Hy-Gō model representation and interactions for (AAQAA)_3_. (Left) Illustration of a native contact between C_α_ atoms and a generic contact between C*β* atoms, along with the corresponding parameters, {*∊*_nc_, *∊*_ab_, *∊*_hp_}, associated with these interactions. (Right) The top two panels present the interaction potentials for the Gō- type interactions as a function of the model parameters. In the top panel, *∊*_db_ = 0.5*∊*_nc_. The bottom panel presents the Weeks-Chandler-Andersen-like potentials employed to model sterics along the peptide backbone.

In addition to the three Gō-type interactions, we also partially employed a standard AA force field, AM-BER99sb,^18^ to model both the steric interactions between all non-hydrogen atoms and also the specific local conformational preferences along the chain. More specifically, the bond, angle, dihedral, and 1–4 interactions of the AA force-field are employed without adjustment. To incorporate generic steric effects, without including specific attractive interactions, we constructed Weeks-Chandler-Andersen potentials^19^ (i.e., purely repulsive potentials) directly from the Lennard-Jones parameters of each pair of atom types in the AA model (bottom panel, Figure 1). For simplicity of implementation, we then fit each of these potentials to an *r*^−12^ functional form. The van der Waals attractions and all electrostatic interactions in the AA force field were not included and water molecules were not explicitly represented. The total interaction potential for the model may be written: *U*_tot_ = *∊*_nc_*U*_nc_ + *∊*_db_*U*_db_ + *∊*_hp_*U*_hp_ + *∊*_bb_*U*_bb_, where the backbone (bb) interaction includes both the intramolecular and steric interactions determined from the AA force field. The first three coefficients represent the only free parameters of the model, while *∊*_bb_ = 1.

The philosophy of this model is that the three Gō- type interactions will *roughly* sample the correct conformational ensemble for short peptides, while atomically-detailed local sterics restrict the model to sample a physically-realistic ensemble of structures. More specifically, the steric interactions ensure that *(i)* conformations that are sterically forbidden in the all-atom (AA) model *are not* sampled and *(ii)* the relevant regions of the Ramachandran plot *are* sampled, while retaining barriers between metastable states. We have previously determined that these characteristics are essential for constructing models with reasonable kinetic properties.^9, 20^ Although the local sterics of the backbone are modeled with near-atomistic resolution, the Hy-Gō model substitutes the full atomistic description of the peptide with a highly coarse-grained representation by modeling the complex combination of dispersion and electrostatic peptide-peptide and peptide-solvent interactions with a limited set of simple interactions between C_*α*_ and C_*β*_ atoms. Note that this is quite distinct from all-atom Gō ^21^ models, which employ atomically-detailed energetic parameters based on the positions of each atom in the native structure.

#### 2. PLUM

As an alternative CG model, we considered PLUM, which also describes the protein backbone with near-atomistic resolution, while representing each amino acid side chain with a single CG site, within an implicit water environment.^6^ In PLUM, the parametrization of local interactions (e.g., sterics) aimed at a qualitative description of Ramachandran maps, while longer-range interactions—hydrogen bond and hydrophobic—aimed at reproducing the folding of a three-helix bundle, without explicit bias toward the native structure.^6^ The model is transferable in that it aims at describing the essential features of a variety of amino-acid sequences, rather than an accurate reproduction of any specific one. After parametrization, it was demonstrated that the PLUM model folds several helical peptides,^6, 22–25^ stabilizes *β*-sheet structures,^6, 26–29^ and is useful for probing the conformational variability of intrinsically disordered proteins.^30^ We also considered 4 minor reparametrizations of the PLUM model:

1. the side chain van der Waals radius is decreased to 90% of its original value.^20^
2. the hydrogen-bonding interaction strength is decreased to 94.5% of its original value.^30^
3. the hydrogren-bonding interaction strength is decreased to 90% of its original value.
4. the side chain interaction interaction strength is decreased to 95% of its original value.

Although PLUM provides a near-atomistic representation of the peptide backbone, the representation is coarsened with respect to the atomically-detailed backbone and side chain sterics of the Hy-Go model. It is important to note that while parametrizations 2-4 change only energetic parameters (i.e., approximately fixed conformational entropy), parametrization 1 significantly changes the conformational entropy of the system by adjusting the steric interactions of the amino acid side chains. In the following, we refer to the latter as the PLUM-ent model.

### B. Simulation Details

In this study, we consider the capped helix forming peptide AC-(AAQAA)_3_-NH_2_, which has been extensively characterized both computationally and experimentally^31^ and is often employed as a reference system for force field optimization. Throughout the manuscript, we refer to the system simply as (AAQAA)_3_.

#### 1. Hy-Gō

CG molecular dynamics simulations of (AAQAA)_3_ with the Hy-Gō model were performed with the Gromacs 4.5.3 simulation suite^32^ in the constant NVT ensemble, while employing the stochastic dynamics algorithm with a friction coefficient γ = (2.0 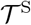)^−1^ and a time step of 1 × 10^−3^ 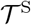. For each model and for each temperature considered, 40 independent simulations were performed with starting conformations varying from full helix to full coil. Each simulation was performed for 100,000 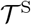, recording the system every 0.5 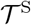. The CG unit of time, 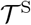, can be determined from the fundamental units of length, mass, and energy of the simulation model, but does not provide any meaningful description of the dynamical processes generated by the model. In this case, 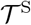 = 1 ps.

#### 2. PLUM

CG simulations of (AAQAA)_3_ with the PLUM force field^6, 24, 33^ were run using the ESPResSO simulation package.^34^ For details of the force field, implementation, and simulation parameters, see Bereau and Deserno.^6^ For each temperature considered, a single canonical simulation was performed for 2,500,000 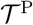 with at timestep of 0.01 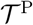, recording the system every 0.5 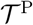, where 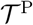 ∼ 0.1 ps. Temperature control was ensured by means of a Langevin thermostat with friction coefficient γ = (1.0 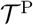)^−1^.

### C. Lifson-Roig Models

According to the Lifson-Roig (LR) formulation,^35, 36^ a peptide system is treated as a 1D-Ising model, which represents the state of each residue as being either helical, h, or coil, c. These simple equilibrium models employ two parameters, w and v, which are related to the free energy of helix propagation and nucleation, respectively. These parameters may be determined directly from simulation data using a Bayesian approach,^31^ and describe the overarching structural characteristics of the underlying ensemble. The average fraction of helical segments, 〈f_h_〉, i.e., propensity of sequential triplets of h states along the peptide chain, is directly determined from {*w, v*}. Although residue- or sequence-specific LR parameters may be determined in order to more faithfully reproduce the helix-coil properties generated from a simulation of a particular peptide system, in this work we determine a single set of {*w, v*} for each simulation model. These simple models are incapable of describing certain features, e.g., end effects, of the underlying systems; however, we utilize the LR parameters only as a characterization tool. For consistent comparison with the experimentally-determined *w* parameter, we determine *w* for each model while setting *v* = 0.05.

### D. Markov State Models

Given a trajectory generated from molecular dynamics simulations, Markov state models (MSMs) attempt to approximate the slow modes of the exact dynamical propagator with a finite transition probability matrix, **T**(τ).^37–39^ This requires a discretization of configuration space, which groups all possible configurations into a manageable set of microstates. Once the microstates are chosen, the number of observed transitions from microstate *i* to *j* at a time separation τ, 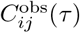, is determined. The matrix of transition counts, **C**^obs^(τ), embodies the dynamics of the simulation trajectory. An estimator for the transition probability matrix is then constructed such that the simulation data is optimally described, while ensuring normalization and detailed balance constraints. The latter constraint applies to any system at equilibrium and alleviates finite-sampling issues. The transition probability matrix is constructed by maximizing the posterior 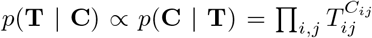, where we applied Bayes’ theorem and a uniform prior distribution.^40^

In the present work, MSMs are built from the (AAQAA)_3_ helix-coil trajectories generated by each CG model. To determine the MSM microstate representation, time-lagged independent component analysis^41^ was performed on the configurational space characterized by the ϕ/ψ dihedral angles of each residue along the peptide backbone. A density clustering algorithm, developed by Sittel *et. al.*,^42^ was then applied to the five “most significant” dimensions in order to determine the number and placement of microstates. We employed the same parameters as used in the original publication for clustering along an all-atom trajectory for Ala_7_. In particular, we used a radius of *R* = 0.1, an energy spacing of 0.1 *k_B_T* for the free energy screening, and an energy spacing of 0.2 *k_B_T* for constructing the network of clusters. A population minimum of 400 and 200 samples per cluster were employed for the Hy-Gō and PLUM models, respectively.

This semi-automated procedure for determining the relevant microstates yields a different number of microstates for each system and for each temperature considered. To estimate the average structural characteristics for a given microstate, we first performed a simple regular space, 2-state clustering along the *ϕ* dihedral angle for each peptide bond. This resulted in a dividing surface approximately at the barrier between *α* and *β* metastable states on the Ramachandran plot. We denoted *α* states as h and *β* states as c and determined an approximate trajectory of the sequence of h/c states. We then compared this trajectory with the trajectory generated from the density clustering and determined the average h/c content for each microstate. These quantities were used to determine the Lifson-Roig parameters, but do not inform the construction of the MSM in any way.

Following the determination of microstates, the simulation trajectories were “cored” using the most probable path analysis^43^ to avoid imperfect definitions of dividing surfaces between microstates. A constant waiting time was used for all microstates and was determined for each system individually in order to ensure that the resulting implied timescales of the model were approximately constant with increasing lag time. This resulted in waiting times between 20 and 100 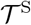(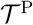) depending on the particular parametrization of the Hy-Gō (PLUM) model and the simulation temperature. Using the cored microstate trajectories, MSMs were generated via a standard maximum-likelihood technique.^40^ For each MSM, the lag time was chosen in the normal way by constructing MSMs for increasing lag times to determine when the resulting set of timescales were converged. Typically, the lag time closely corresponded to the waiting time used in the coring analysis and did not exceed three times this number. Figure 2(a) presents a representative implied timescale plot. To validate the resulting models, we performed the standard Chapman-Kolmogorov tests in order to compare the probability decay from various metastable states estimated from the simulation and predicted from the MSM. Figure 2(b) presents a representative Chapman-Kolmogorov result for the decay of probability from the helix state as a function of time. MSM construction and analysis were performed using the pyEmma package.^44^

**FIG. 2.**
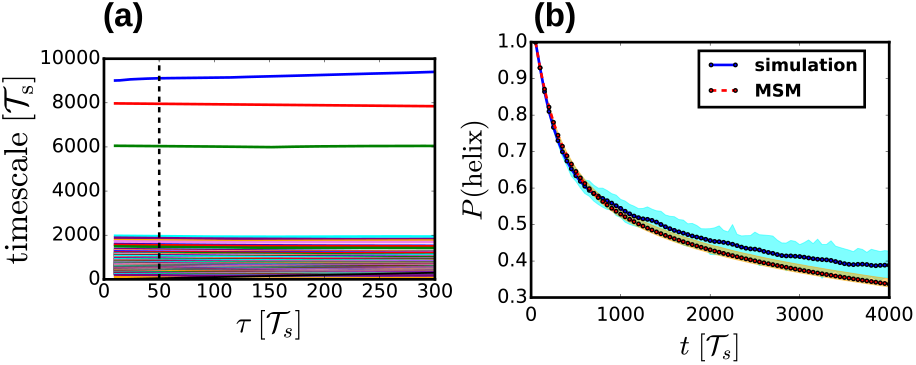
(**a**) Representative implied timescale test. Each line represents a different characteristic timescale of the Markov state model as a function of lag time. The vertical dashed line denotes the lag time chosen in this case. (**b**) Representative Chapman-Kolmogorov test. The “helix” state corresponds to the microstate for each model with the highest helical content (see Methods section for details). The blue solid curve presents the probability decay of the helix state, as determined directly from the simulation trajectory, while the red dashed curve presents the same quantity determined from the MSM. The transparent cyan and orange regions denote the error bars for each quantity.

## III. RESULTS AND DISCUSSION

In this study, we investigate the relationship between structural and kinetic properties of helix-coil transition networks generated by microscopic simulation models. As a structural-characterization tool, we construct Lifson-Roig (LR) models from each simulation trajectory. Although the kinetics of helix-coil transitions are often interpreted in terms of a kinetic extension of the Ising model,^45, 46^ the precise impact of the model’s assumptions on the fine details of the resulting kinetic network is not well understood. For example, we recently demonstrated that the topological details of the kinetic networks generated by various simulations models for Ala_7_ may vary drastically, while generating nearly identical structural properties.^13^ Here, we construct Markov state models (MSMs) directly from the simulation trajectories, allowing a more complex relationship between the LR parameters and the kinetic properties of the system.

As a model system, we investigate the AC-(AAQAA)_3_-NH_2_ peptide, which contains 15 peptide bonds. From the LR point of view, there are 2^15^ states, determined by enumerating the various sequences of helical (h) and coil (c) states along the peptide backbone. However, this representation is inappropriate for constructing MSMs from the simulation trajectories. Instead, we systematically determine coarser, representative microstates, which correspond to collections of the more detailed configurations of the system. See the Methods section for a detailed description of microstate determination. For reference, we employ data from previous analysis of the (AAQAA)_3_ system.^31^ In particular, we characterize the ensembles generated by CG simulation models with respect to those obtained from NMR experiments and from simulations of the ff03* all-atom (AA) model.

We consider two distinct types of CG models in this study, as described in greater detail in the Methods section. First, we employ a relatively simple, native-biased CG model. The hybrid Gō (Hy-Gō) model, is a flavored-Go model,^14–16^ with 3 Gō-type interactions and also physics-based interactions in the form of sterics and torsional preferences along the backbone.^13^ We constructed 15 distinct Hy-Gō parametrizations by varying the Gō-type interactions while keeping the steric interactions fixed (i.e., the conformational entropy of the models remains approximately unchanged). We also considered the transferable PLUM model^6^ along with 4 minor reparametrizations. 3 of the reparametrizations also corresponded to changing only the energetic parameters, while 1 reparametrization changed the side chain bead size and, thus, the conformational entropy with respect to the other PLUM models. We simulated each model over a range of temperatures to assess their thermodynamic properties (i.e., the temperature dependence of structural quantities). The energy scales of the models were aligned by shifting the temperature scale such that the average fraction of helical segments (i.e., 3 consecutive helical states), 〈*f*_h_〉, at the reference temperature, *T*\*, matched the experimental value at 300 K. For each model and each temperature considered, we determined the two LR parameters, {*w, v*}, and constructed an MSM directly from the simulation data.

### A. Properties of the (AAQAA)_3_ helix-coil transition

We characterized the overall structure of each ensemble in terms of the LR free energy of helix extension, *w*, and the average fraction of helical segments, 〈*f*_h_〉. Figure 3(a) and (b) present these two quantities, respectively, as a function of temperature for the 15 Hy-Gō models (colored curves with circle markers) and 5 PLUM models (gray-scale curves). The original PLUM model and 3 energetic reparametrizations are denoted with square markers, while the steric reparametrization, named “PLUM-ent”, is denoted with an X marker. From the temperature dependence of *w*, we fit a thermodynamic model to quantify the *T*-independent enthalpy and entropy of helix extension. In particular, we assume *k_B_T* ln *w(T)* ∼ Δ*H*_hb_ – *T*Δ*S*_hb_, neglecting the heat capacity contributions to the free energy and considering a relatively small range in *T*. Δ*H*_hb_ corresponds to the slope of the curves in Figure 3(a) and is a simple measure of the cooperativity of the transition. For consistency with the CG models, we fit the experimental *w* values from a similarly small temperature range, resulting in a slightly inflated experimental Δ*H*_hb_ compared with that reported in previous work.^31^

**FIG. 3.**
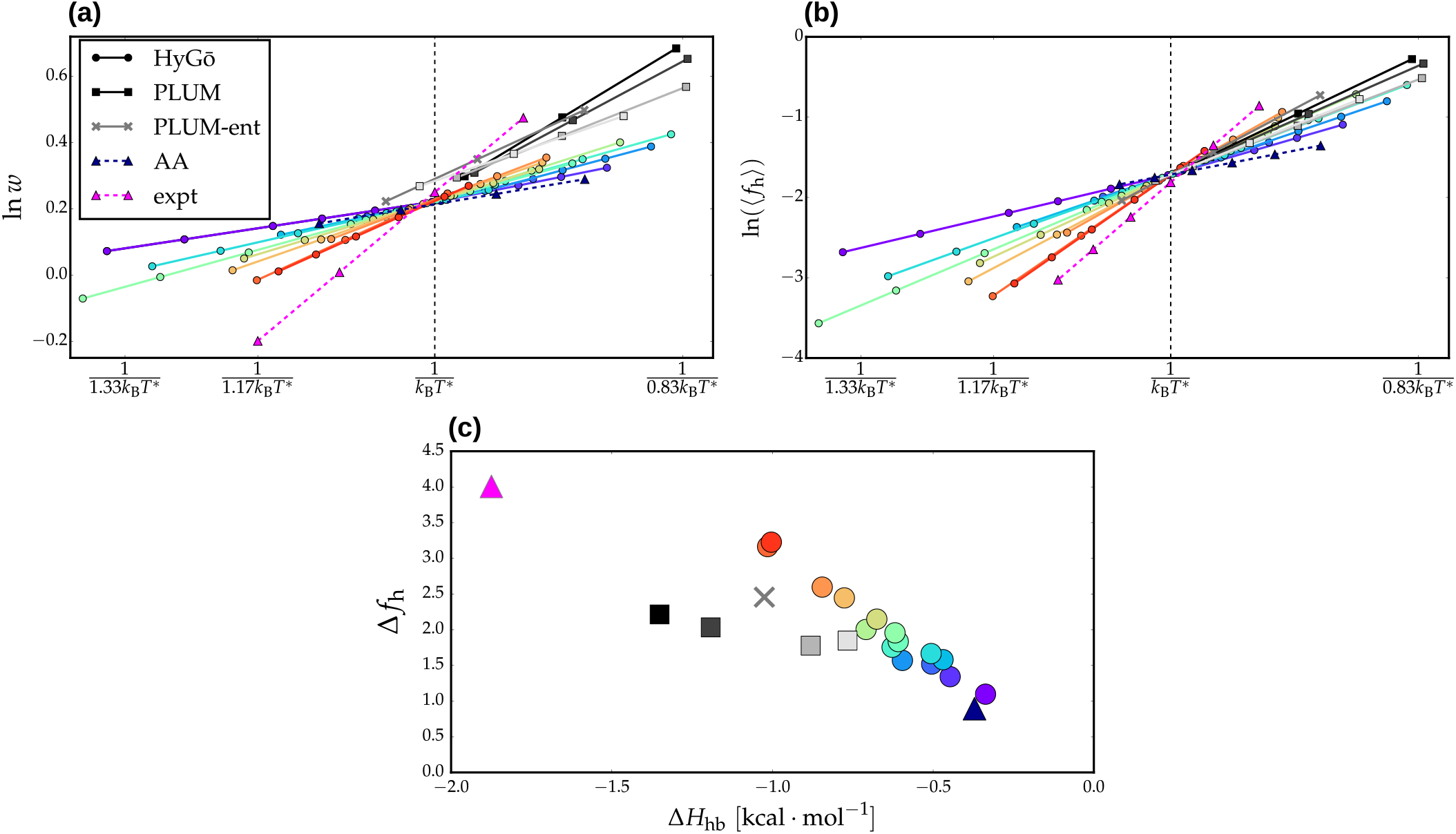
Temperature dependence of the Lifson-Roig helix propagation parameter, *w* (**a**), and the average fraction of helical segments, 〈*f*_h_〉 (**b**). The curves presented are a linear fit of the raw data, (**c**) The slope of ln〈*f*_h_〈(1/*T*), denoted Δ*f*_h_, versus the enthalphy of helix extension (determined as the slope of ln *w*(1/*T*)). Data from the Hy-Gō models are denoted with circle markers and are colored according to their relative cooperativity (as determined by Δ*f*_h_). The results from the PLUM models are denoted with square markers and are colored in gray-scale according to Δ*H*_hb_. The experimental and all-atom results taken from Best and Hummer^31^ are denoted with triangle magenta and dark blue markers, respectively. Note that the models are aligned by shifting the temperature such that all models achieve 〈*f*_h_)^expt^ at *T*\*.

Figure 3(c) presents the slope of ln〈*f*_h_〉(1/*T*), denoted simply Δ*g*_h_, versus Δ*H*_hb_ for each model. Recall that within the LR formulation, both the helix extension parameter, *w*, and the helix nucleation parameter, *v*, contribute to determining the average fraction of helical segments, 〈*f*_h_〉. Since it is typically assumed that v is dominated by entropic contributions,^36^ one may expect a linear correlation between Δ*f*_h_ and Δ*H*_hb_ for models with similar conformational entropy (i.e., models that differ only by their energetic parameters). (Note that although we determine LR models with fixed *v*, the corresponding entropic effects are effectively folded into *w* to generate the appropriate 〈*f*_h_〉). Indeed, Figure 3(c) demonstrates that both the Hy-Gō and PLUM models (except the PLUM-ent model, gray X marker) generate individual linear correlations between Δ*f*_h_ and Δ*H*_hb_. This is consistent with our previous results for Ala_7_.^13^ For models with differing conformational entropy, we expect, in general, distinct trends between Δ*f*_h_ and Δ*H*_hb_, as long as entropy is playing a significant role in determining 〈*f*_h_〉. This is clearly the case for the Hy-Gō and PLUM models for (AAQAA)_3_ (Figure 3(c)). In our previous investigation of Ala_7_,^13^ we found a consistent linear correlation for both model types. This could be because either 1. entropy does not play a significant role in determining 〈*f*_h_〉 or 2. the PLUM models considered happened to fall on the intersection of the two linear trends. The implication of conformational entropy as the dominant factor in the difference between the two linear correlation trends in Figure 3(c) is strongly supported by the PLUM-ent model, which represents a change in the conformational entropy via adjustment of steric interactions. For this model, the van der Waals radius of the side chain was adjusted to better represent the Ramachandran regions sampled by a higher-resolution model. This adjustment results in a Δ*f*_h_/Δ*H*_hb_ trend in closer agreement with the Fly-Go models, which employ detailed steric interactions that faithfully describe the conformational entropy of the peptide backbone. Figure 3(c) demonstrates that the slope of the Δ*f*_h_/Δ*H*_hb_ correlation for the various parametrizations of the Fly-Gō model is consistent with both experimental and AA results.^31^

### B. Validation of structural-kinetic relationships for (AAQAA)_3_

Our previous work identified a robust relationship between structural and kinetic properties generated by distinct models for Ala_7_.^13^ In particular, it was demonstrated that the average fraction of helical residues, 〈*N*_h_〉, and the average fraction of helical segments, 〈*f*_h_〉, determined the ratio of rates for characteristic nucleation and elongation processes. The calculation of nucleation and elongation rates for Ala_7_ depended upon the identification of particular microstates representing nucleation states of the peptide. Here, the situation is complicated by our heterogeneous microstate representation for the (AAQAA)_3_ ensembles. For this reason, we first considered coarser processes—global folding and unfolding of the peptide. We define the timescales of folding, *t*_fol_, and unfolding, *t*_unf_, as the mean first passage times from the coil ensemble to the full helix state and vice versa, respectively. The coil ensemble is comprised of all microstates whose average structure contains no consecutive set of three residues with greater than 50% helicity (according to a naive dividing surface definition of the helical state of a single residue, see Methods section for details). Subsequently, we also considered faster processes corresponding to the waiting time that each residue, *i*, spends in a helical state before transitioning to the coil state, 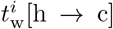, and vice versa, 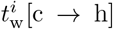. These waiting times were calculated using the 2-state dividing surface along the ramachandran plot, as described in the Methods section. Note that since the CG unit of time does not correspond to a physical time, there is always an arbitrary speed-up associated with dynamical properties generated by CG models. For this reason, we only consider ratios of timescales, to effectively account for this speed-up.

Figure 4(ai) presents the temperature dependence of the ratio of folding to unfolding timescales. Remarkably, the Hy-Gō models generate nearly identical folding versus unfolding timescales at the reference temperature *T\**, regardless of the particular parametrization. The PLUM models with similar conformational entropies also demonstrate similar folding to unfolding ratios, although the variance is somewhat higher than the Hy-Gō models. Recall that the energy scales of the models are aligned by shifting the temperature such that all models have the same value of 〈*f*_h_〉 at *T\**. Thus, similar to our previous results for nucleation/elongation timescales, the folding/unfolding timescales are largely determined by the average fraction of helical segments in a consistent manner over all parametrizations of a given model type. However, in this case, there is a shift in the timescale ratio at the reference temperature depending on the details of the model representation. This is supported by the fact that the PLUM-ent model (Figure 3(c)) displays kinetics at the reference temperature which are closer in line to the Hy-Gō models. The validity of the structural-kinetic relationship is further demonstrated in Figure 4(aii), which presents the slope of the curves in panel (ai) versus the enthalpy of helix elongation, Δ*H*_hb_. Interestingly, the temperature-dependence of the timescale ratio is dictated by Δ*H*_hb_, determined from the LR model, rather than the slope of the actual helical content, Δ*f*_h_, albeit these two quantities largely coincide within a model class. It is noteworthy that this particular structural-kinetic relationship is one that is well-known—the ratio of folding to unfolding timescales quantifies the free energy difference between the folded and the unfolded state for a 2-state folder.^47^

**FIG. 4.**
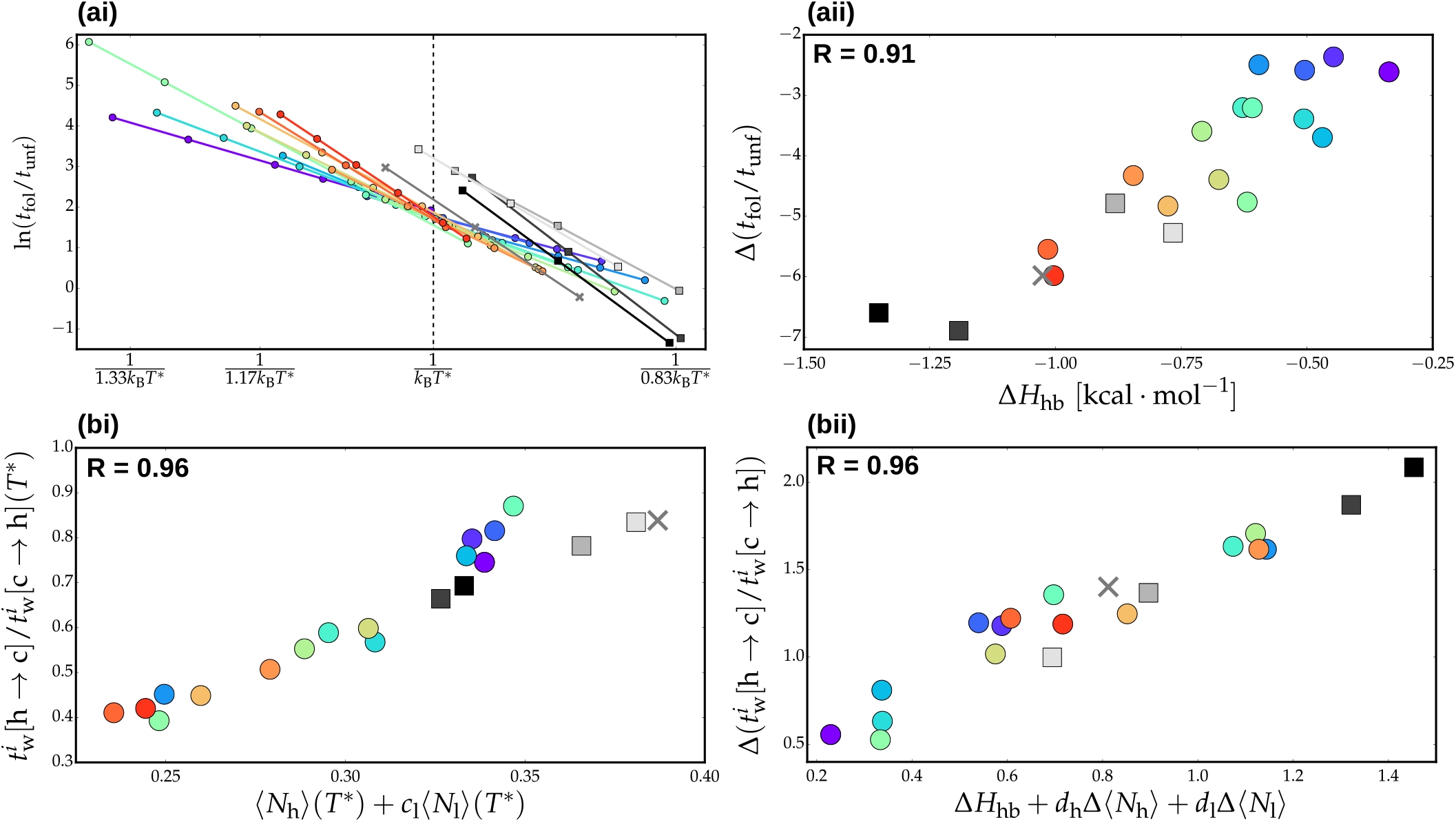
(**ai**) Temperature dependence of the ratio of folding to unfolding timescales, *t*_fol_/*t*_unf_. (**aii**) Slope of the linear fit of ln(*t*_fol_/*t*_unf_) as a function of inverse temperature versus the enthalphy of helix extension, Δ*H*_hb_ (determined as the slope of ln *w*(1/*T*)). (**bi**) Ratio of waiting time in the helix state, 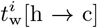, to waiting time in the coil state, 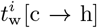, for residue position *i* as a function of the average fraction of helical residues, 〈*N*_h_〉, and the average fraction of lone helices, 〈*N*_1_〉, at *T*\*. The free parameter, *c_l_*, was determined by a fit over all models to maximize the pearson correlation coefficient, R, between the plotted quantities. Here, *i* corresponds to the terminal ends of the peptide (not including capping groups), (**bii**) Slope of the linear fit of 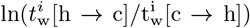 as a function of inverse temperature versus Δ*H*_hb_, the slope of the linear fit of 〈*N*_h_〉 as a function of inverse temperature, and the slope of the linear fit of 〈*N*_l_〉 as a function of inverse temperature. The free parameters, *d_h_* and *d_l_*, were determined by a fit over all models to maximize the pearson correlation coefficient, R, between the plotted quantities.

Panel (b) of Figure 4 presents similar results to panel (a), but for the ratio of waiting time in the helical (h) state to waiting time in the coil (c) state for a particular residue *i*. Here, *i* represents a position on the helix and an average is performed over residues in that position from either end of the helix. In particular, panel (b) presents results from the terminal ends of the helix (excluding capping groups), which corresponds to the fastest h to c transitions along the peptide chain. However, equivalent results were obtained for each position along the peptide. Unlike the folding to unfolding ratios, the various models generate distinct waiting time ratios at *T*\*. However, panel (bi) demonstrates that the differences in the ratios is determined by structural features of the ensemble at T*, namely, the average fraction of helical residues, 〈*N*_h_〉, and the average fraction of lone helices, 〈*N*_l_〉. In other words, while the global folding to unfolding ratio is determined only by the average fraction of helical segments, the consistency of local kinetics requires further alignment of the structural ensemble. Correspondingly, panel (bii) demonstrates that the temperature dependence of the ratio of waiting times depends not only on Δ*H*_hb_ but also the temperature dependence of 〈*N*_h_〉 and 〈*N*_l_〉.

### C. Thermodynamics and transition network topology

Our previous analysis of Ala_7_ suggested that topological features of the transition network at a single reference temperature may be capable of determining the thermodynamics (i.e., temperature dependence) of the model, without explicitly taking into account simulations at various temperatures.^13^ We found that the distribution of paths from full helix to full coil states in the network provided a good characterization of the topology of the network. To quantify this distribution, we employed the conditional path entropy,^48^ *H*_sd|u_, which characterizes the average degree of randomness for paths from *s* to *d* passing through a particular intermediate state, *u*. Our hypothesis was that models with more directed transitions to the helix state, i.e., lower average conditional path entropy in the folding direction, would undergo a faster change in helical content with decreasing temperature (i.e., higher cooperativity). Our results demonstrated that this naive picture was partially valid, although a more complex relationship between the topology and cooperativity was necessary for a quantitative description.^13^

Figure 5 presents the conditional path entropy in the folding direction, averaged over all intermediate states, versus Δ*H*_hb_ for each model. Note that the PLUM-ent model was left out because its microstate representation contained too few states to produce meaningful results with this analysis. It is also important to note that the models were not simulated exactly at *T*\*, introducing errors into any legitimate trend. For numerical convenience, we weighted each of the features by the fractional flux passing through each state. We also chose to normalize the path entropies by the total number of paths for each network. Similar to our previous results, 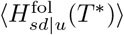 provides a rough description of cooperativity of the model. Extrapolating to the experimental value of Δ*H*_hb_ (∼1.9 kcal · mol^−1^), Figure 5 implies that the experimental helix-coil transition would have a very low conditional path entropy and, thus, undergoes an incredibly directed transition from helix to coil state. This is in line with the conventional view of a 2-state transition, where there is an exponential suppression of intermediate states.^23^

**FIG. 5.**
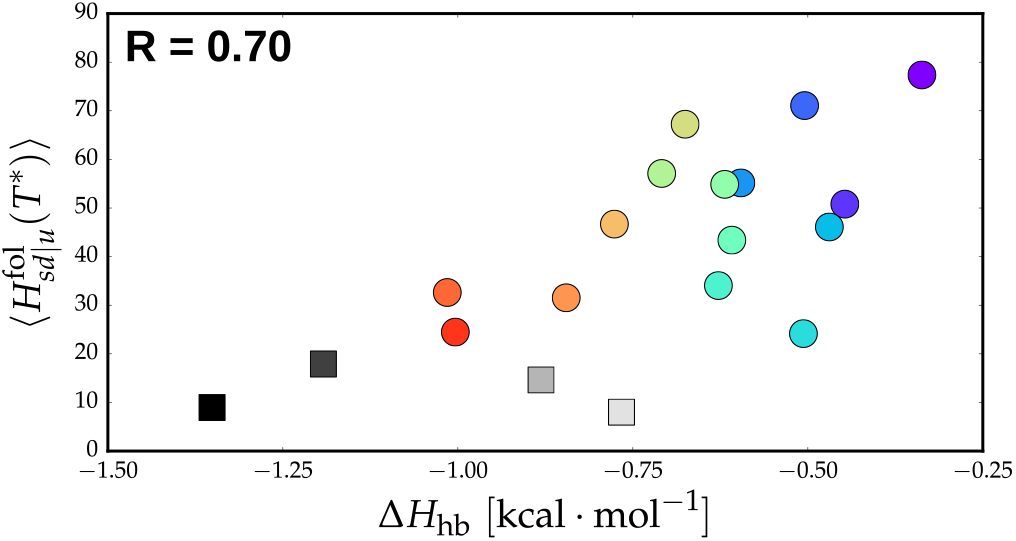
Average conditional path entropy in the folding direction versus the enthalpy of helix extension, Δ*H*_hb_ (determined as the slope of ln *w*(1/*T*)).

## IV. CONCLUSIONS

We recently demonstrated structural-kinetic relationships for peptide secondary structure formation that emerge if a CG model incorporates certain essential physics.^13^ In this work, we further validate these structural-kinetic relationships for a longer peptide, where the representation of configuration space in terms of the full enumeration of sequences of helical/coil states along the peptide backbone is impractical. Furthermore, the role of conformational entropy in the determination of these relationships was clarified by distinguishing between models with fixed or varying steric interactions. More specifically, a change in the conformational entropy, achieved by either changing the model representation or by explicitly adjusting the steric interactions, results in a shift in the global, long timescale kinetic observables at the reference temperature. However, this does not affect the temperature dependence of the kinetic properties, which vary consistently with the enthalpy of helix extension. Additionally, local kinetic properties are determined by structural features of the ensemble, without dependence on the model representation. The semiautomated construction of Markov state models provides a network picture of dynamics, allowing at least the partial characterization of temperature-dependent quantities from a single reference temperature. In combination with our previous work,^9, 13, 20^ the results presented here provide a potential strategy for ensuring kinetic accuracy in CG models through the matching of particular structural quantities. This approach requires further identification and validation of structural-kinetic relationships for tertiary structure formation, and may be especially useful for providing structural interpretations for kinetic protein experiments.

## ACKNOWLEDGEMENTS

We thank Marius Bause and Alessia Centi for critical reading of the manuscript. This work was funded through a postdoctoral fellowship from the Alexander von Humboldt foundation (J.F.R.) and an Emmy Noether fellowship from the German Research Foundation (T.B.).

